# A data-informed approach for engineering in-vitro experiment design to decipher key features of invasive breast cancer cell phenotypes

**DOI:** 10.1101/2025.03.21.643499

**Authors:** Lekha Shah, Valentina Breschi, Annalisa Tirella

## Abstract

The intrinsic complexity of biological processes often hides the role of dynamic microenvironmental cues in the development of pathological states. The use of micro-physiological systems (MPS) offers new technological platforms designed to model the dynamics of tissue-specific microenvironments *in vitro* and to holistically understand healthy and pathological states. In our previous works, we reported on engineering breast critical tumor microenvironment features, including matrix stiffness, pH, and fluid flow, and use the MPSs to study breast cancer cells phenotypes. By studying different microenvironments mimicking normal and tumor breast tissues, we obtained high-dimensional data using two distinctive human breast cell lines (i.e., MDA-MB231, MCF-7) investigating biomarkers commonly used in cancer *in vitro* models as cell proliferation, epithelial-to-mesenchymal transition (EMT), and breast cancer stem cell markers (B-CSC). We herein report on a new approach used to explore the complexity of MPSs and the high dimensional datasets: we introduce an innovative machine learning (ML) based platform employing unsupervised k-means clustering and feature extraction to identify key markers that differentiated invasive from non-invasive breast cell phenotypes. This novel data-driven approach streamlines experimental design and emphasizes the translational potential of integrating MPS-derived insights with ML to refine prognostic tools and personalize therapeutic strategies.

**Graphical Abstract:** 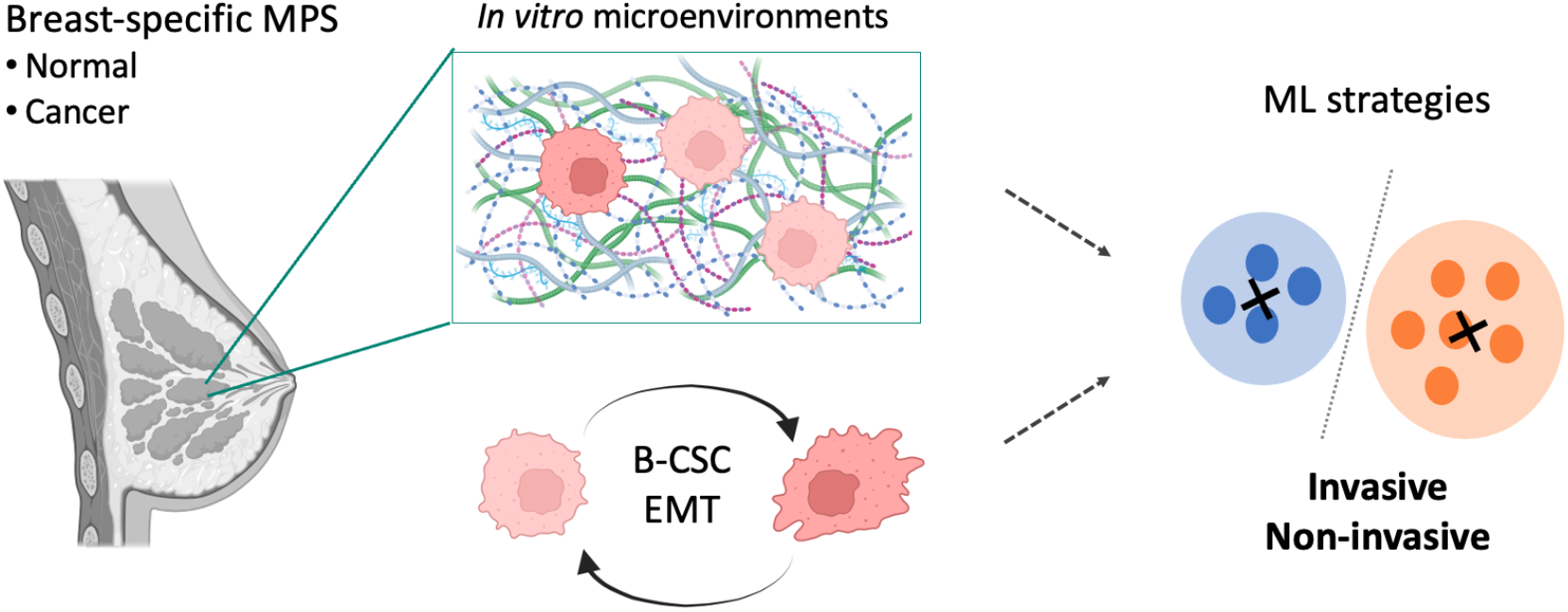

## 1. Introduction

Breast cancer is a significant cause of cancer-related deaths among women, with recent statistics stating 2.3 million cases diagnosed globally in 2022 and an estimated 660,000 deaths ^1^. Most of these deaths are related either to metastatic disease at initial presentation (Stage III and above) or to recurrence at a later stage ^2^. In particular, recurrence occurs in around 20% of breast cancer cases, with the time to recurrence varying from as low as 3 months to as long as 32 years after the initial diagnosis ^3,4^. In this scenario, health technologies are urged to tackle issues such as prevention, precision diagnostics, personalized medicine, and disease management.

In the specific case of breast cancer, improved information on a patient’s risk of recurrence and estimated time to recurrence become very important to guide appropriate and personalized adjuvant therapy, with the possibility to effectively plan future screening programs. In fact, the risk of breast cancer recurrence and metastasis has already been linked clinically to different factors such as molecular subtypes, grade, tumor node status, size of initial tumor ^3^ and patient’s status (age and menopause) ^5^. In line with these results, improvements toward a more patient-centered approach in prognostic tests for different stages and identification of appropriate treatment have been implemented ^6,7^. For example, among the molecular subtypes, triple negative breast cancer (TNBC) and Her2+ subtypes have more recurrence risk and lower average recurrence time (26 months) as compared to luminal A and luminal B subtypes (56 months) ^3^ Other tools, such as Nottingham prognostic index take into consideration only histological factors ^8^, whereas newer ones such as MammaPrint, Oncotype DX, Mammostrat, Prosigna, EndoPredict, IHC4 are more patient-specific with relevant gene expression and biomarker analysis ^6^. However, all these tests represent a snapshot of each patient’s condition and lack information on the time-variance of tumor progression, metastatic onsets and possible metastatic dissemination. It is known that at the time of diagnosis of breast cancer, an estimated 75% of initial tumors are already disseminated to a distant site ^9^. These micro-metastatic sites persist in a state of dormancy within the body until the onset of full metastatic expansion eventually occurs, which is known to correlate with tumor cell phenotypes and tumor microenvironment (TME). Therefore, identifying which features of breast tumor ecosystem trigger onset of tumor progression and dissemination is key enabler for precise prognosis and identification of efficacious treatments.

Preclinical animal models, like patient-derived xenografts are considered the most promising *in vivo* model to study spatial structure, heterogeneity and genomic features of human cancers for identification of effective treatments ^10^. However, these approaches are prone to ethical concerns, while also being time-consuming and costly in developing patient-specific and patient-centered models ^10,11^. In addition, such *in vivo* models often do not replicate the rate of dissemination and metastasis, due to a poor representation of human-specific metastatic sites.

Preclinical *in vitro* models, often considered too simplistic, can now integrate technological advancements, such as new biomaterials in microfluidic technologies, and together with digital data and tools are considered key enablers for a more precise and sustainable patient-centered solution to tackle human diseases. The main advantage of engineered *in vitro* models is their ability to mimic multiple aspects of tissue-specific dynamics whilst precisely controlling the ecosystem, spanning from the selection of cell types to the physical properties of the extracellular matrix (ECM). Thanks to these technological advancements and integrations, engineering tissue-specific key features in *in vitro* models raises debate over whether these or *in vivo* models serve as better models for understanding tumor dynamics and metastatic onsets ^12,13^.

Micro-physiological systems (MPS) are microscale three-dimensional (3D) *in vitro* cell culture platforms modelling the functional features of tissues by exposing cells to a combination of physical (e.g., temperature, pH, oxygen), biochemical, electrical, mechanical (e.g., flow, stretch), structural, and/or morphological conditions to replicate healthy or diseased tissue/organ functions ^14^. In this perspective, we engineered 3D *in vitro* breast cancer models for mechanistic discovery and tailoring TMEs with reproducible and customizable physico-chemical properties to match the breast-specific microenvironment ^15^. Such breast-specific MPS enabled independent modulation of physico-chemical properties of the ECM (i.e., mechanical properties, density and composition), environmental pH and fluid flow (**Figure 1**). Alginate-based hydrogels were used to tailor compressive moduli in the range of 2-10 kPa, as reported to be clinically relevant in the context of breast cancer matrix stiffness ^16,17^. The combination of mechanically tunable alginate hydrogels with ECM-derived materials (i.e., gelatin) returned interpenetrated hydrogel networks with controlled density, directly linked to clinically relevant markers in breast cancer diagnosis, that can be easily integrated in a fluidic chamber to study cell-matrix interaction under fluid flow mediated force transmission. Based on our experience in analyzing both individual and combinatorial effects of breast-specific TMEs on two selected human breast cancer cell lines (i.e., the highly invasive/recurrent TNBC cell line MDA-MB 231, the less invasive/low recurrence luminal A cell line MCF-7) by measuring their proliferation, breast cancer stem cell (B-CSC) population and epithelial-to-mesenchymal transition (EMT) ^18,19^, we correlated that TME key features can modulate malignant phenotypic transitions such as EMT/MET, and also have a far-reaching impact by increasing invasive potential and bone metastatic osteolysis *in vitro* ^15,20^. This confirms that breast-specific MPS could inform not only on the current molecular state of the tumor but also on its potential to change into malignant phenotypes, given the relevant micro-environmental conditions.

**Figure 1.**
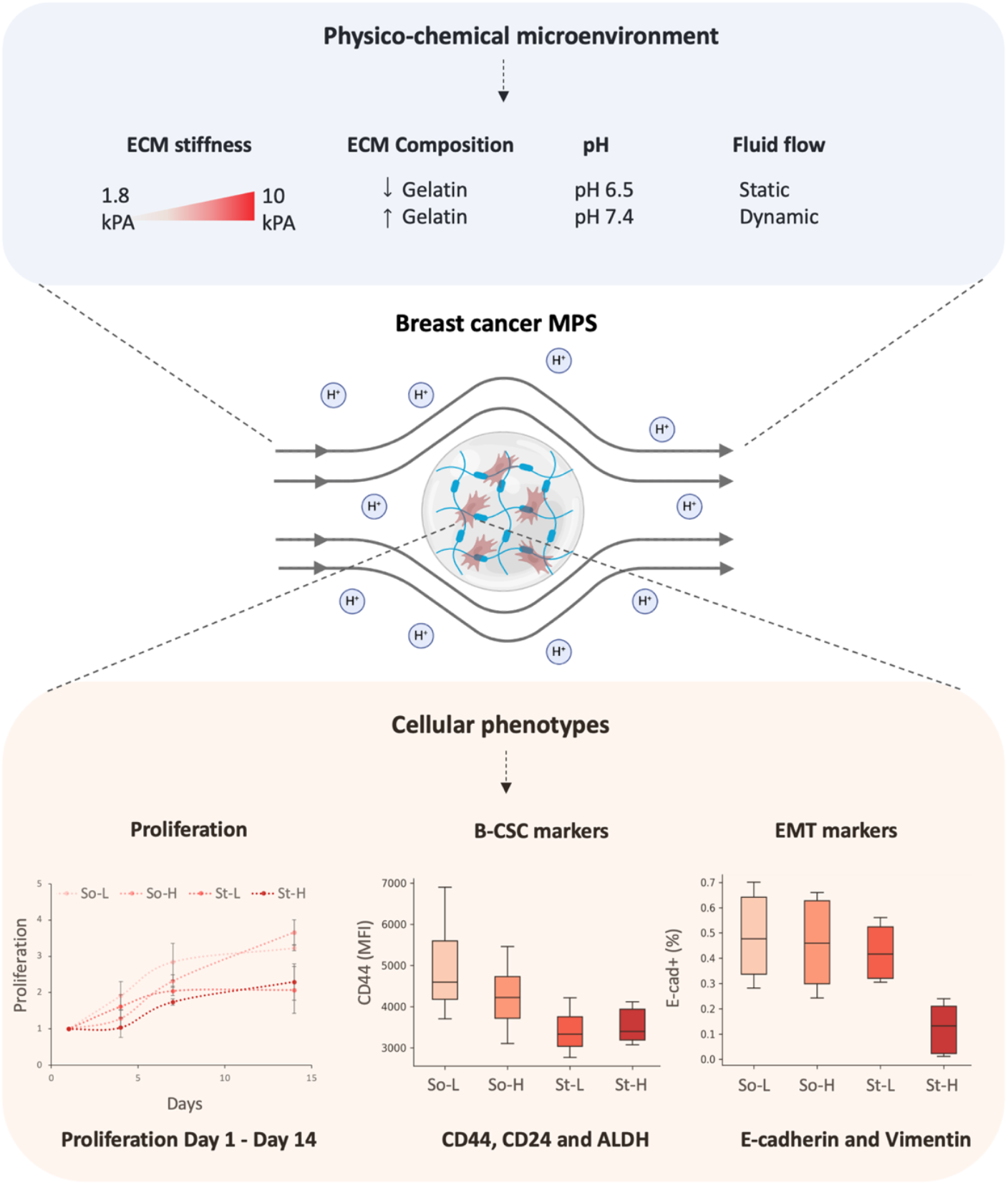
Experimental design representing breast-specific MPS and biological readouts. Schematic representation of human breast cancer cells (i.e., MDA-MB 231, MCF-7) encapsulated in hydrogels with tailored physical properties (i.e., Young’s modulus, composition, density) cultured in controlled culture system (i.e., pH, perfusion). Cellular phenotypes, such as proliferation (Alamar blue assay), flow cytometry analysis of B-CSC marker expression (CD44, CD24, ALDH) and EMT marker expression (E-cadherin and Vimentin), were monitored and recorded over time. Example of obtained data (bottom), cell proliferation curves, CD44 median fluorescence intensity, and E-cadherin+ (%) data of MCF-7 cells cultured in four different hydrogels (So-L, So-H, St-L and St-H) at pH 7.4 and static conditions.

Whilst designed to offer a better understanding of cell-matrix interactions and linked biological processes, MPS models often require a rather long execution phase to first validate physico-chemical features of the TME, followed by time-consuming experimental campaigns with time-points decided based on previous studies, and not tailored to follow the dynamic of the specific model. Experimental campaigns often collect high dimensional datasets, which increase proportionally to the number of controlled variables of the MPSs used, often resulting in readouts that could be difficult to fully interpret and understand. Indeed, this complexity hinders a clear understanding of each variable’s role, for example in guiding cell phenotype behaviors, ultimately preventing the identification of parameters of interest to distinguish cellular phenotypes and, therefore, to inform on disease progression (e.g., invasion, distal metastasis). Techniques drawn from statistics and machine-learning (ML) can be used to disentangle retrieved information, setting the ground for sustainable design approaches that emphasize relevant TME features influencing trial’s outcome and identify a limited pool of key defining variables. As a consequence, ML not only provides a set of approaches to better understand patterns in cells’ behaviors linked to the TME features, but it can empower future *in vitro* experiment design.

Based on these observations, we herein collected all experimental datasets from previous studies and used a ML approach to better understand the impact of the different engineered features of the breast cancer *in vitro* models and the relevance of the measured outcomes in shaping and describing specific cellular and biological readouts, with the final goal to achieve a sustainable-by-design experimental campaign and simplify the analysis of trials’ readouts towards a patient-centered predictive model (**Figure 1**). However, unveiling these features is all but a trivial task. Indeed, data from preliminary *in vitro* experiments are often scarce with respect to the data sizes required for complex ML techniques, which are knowingly data-hungry, especially when the readouts require time-consuming manual sample preparation. To overcome these limitations, we propose to leverage (simple) unsupervised ML strategies and feature importance techniques to learn the engineered variables that allow for distinguishing cell phenotypes based on data gathered in our already described breast cancer *in vitro* models ^15^. The resulting combination of breast-specific MPS and (simple) ML models is used to investigate the effect of TME on invasive potential and/or recurrence risk, toward achieving the ultimate goals of this new study: 1) uncovering the biomarkers that should be monitored to detect invasive breast cancer cell phenotypes over non-invasive ones for an in-depth characterization; and 2) understanding whether these biomarkers depend on the properties of the MPS model. To the best of our knowledge, this approach has not been reported previously, providing a unique and innovative platform for studying such critical cellular processes in a highly physiological and pathological context.

## 2. Materials and Methods

### 2.1 Data collection

In this study, we included 12 different tissue-specific microenvironments (**Table 1**) and used with two different human breast cancer cell lines and obtained 24 breast-specific MPSs; **Table 2** reports the 16 different cellular responses/outputs measured for each condition and cell line used.

**Table 1.** Breast-specific MPSs (12) and corresponding physical features used to model healthy and tumoral states. Alginate-based hydrogels were classified based on their mechanical properties (E, Young’s modulus) and density (gelatin content). Inclusion of cell-laden hydrogels in bioreactor was used to transmit forces (dynamic) and compared to traditional cell culture (static). The pH of cell culture medium was buffered to values reported for healthy (pH 7.4.) or tumor (pH 6.5) tissues.

| MPS | Hydrogel<br>($E$, kPa) | Alginate<br>(% w/v) | Gelatin<br>(% w/v) | pHe | Fluid mediated<br>flow/ force |
| --- | --- | --- | --- | --- | --- |
| 1 | So-L<br>($1.8 \pm 0.2$) | 1.5 | 1 | 7.4 | Static |
| 2 |  |  |  | 6.5 | Static |
| 3 |  |  |  | 7.4 | Dynamic |
| 4 | So-H<br>($2.4 \pm 0.1$) | 1.5 | 3 | 7.4 | Static |
| 5 |  |  |  | 6.5 | Static |
| 6 |  |  |  | 7.4 | Dynamic |
| 7 | St-L<br>($6.1 \pm 0.2$) | 3 | 1 | 7.4 | Static |
| 8 |  |  |  | 6.5 | Static |
| 9 |  |  |  | 7.4 | Dynamic |
| 10 | St-H<br>($10.1 \pm 0.5$) | 3 | 3 | 7.4 | Static |
| 11 |  |  |  | 6.5 | Static |
| 12 |  |  |  | 7.4 | Dynamic |

So = soft hydrogel ( $E < 4$ kPa, normal); St = stiff hydrogel ( $E > 5$ kPa, tumour); L = low gelatin content; H = high gelatin content.

**Table 2.** Experimental data used as output of tested MPS models (n=12; Table 1) for each human breast cell line (m=2; i.e., MCF-7, MDA-MB231).

| Experimental method | Measured data |
| --- | --- |
| Cell Proliferation<br>*Alamar blue assay* | Metabolic activity in time (time points: day 1, day 4, day 7, day 14) |
| Breast cancer stem cell markers (B-CSC)<br>*Flow cytometry analysis* | CD44 (MFI), CD24 (MFI), CD44v6 (MFI), CD44+ (%), CD24+ (%), CD44+/CD24- (%), CD44v6+ (%), ALDH+ (%) |
| Epithelial-to-mesenchymal markers<br>*Flow cytometry analysis* | Vimentin (MFI), E-cadherin (MFI), Vimentin+ (%), E-cadherin+ (%) |

#### 2.1.1 Engineered 3D breast in vitro models

##### Human breast cells

Human breast adenocarcinoma cell lines MCF7 and MDA-MB-231 were selected and authenticated by European Collection of Authenticated Cell Cultures (ECACC, operated by Public Health England) prior use. Cells were routinely cultured in complete medium and were discarded after they reached passage 25 for MDA-MB-231 and passage 50 for MCF-7. In specific, two breast cancer cell lines were chosen to represent opposite ends of the breast ‘invasive potential’ spectrum: the highly invasive triple negative cancer cell line (MDA-MB 231) and less invasive luminal type (MCF-7).

##### 3D *in vitro* models

Alginate-based hydrogels characterized in our previous study ^15^ were selected based on their elastic modulus (soft vs stiff), density and gelatin content (low vs high) and to mimic characteristics of normal (soft, low) and cancer breast tissue (**Table 1**). Hydrogels were designed to target Young’s modulus values (also referred to as stiffness) from 1 kPa (normal breast tissue) to 10 kPa (tumor breast tissue); low and high gelatin concentration to mimic either normal breast tissue or collagenous dense tumor tissue. For 3D cell culture studies, spherical hydrogels beads were prepared gently re-suspending cells in alginate precursor solutions (aq.) ensuring a homogeneous single cell suspension and at a concentration of 10^6^ cells/mL. Then single droplets of cells suspended in alginate were ejected from a 25G needle in a sterile crosslinking solution allowing gelation for 10 min at room temperature (RT).

##### Environmental pH

Different media composition was used to model pH variations and tailor a normal (i.e. pH 7.4) and a cancer (i.e. pH 6.5) microenvironment. DMEM buffered cell culture media were prepared with HEPES-PIPES ^15^. To ensure maintenance of the target environmental pH, cell culture media was changed every two days, with cells cultured in standard conditions (37°C, 5% CO_2_).

##### Intra-tumoral pressure: bioreactor and force transmission

To model force transmission as in TME, interstitial fluid flow of bioreactor was used to transmit mechanical forces to cells: the Quasi vivo QV500 system (Kirkstall, UK) equipped with the Watson-Marlow 202U peristaltic pump was used as reported in ^15^ and setting a constant flow rate of 500 μL/min.

#### 2.1.2 Cell viability

Alamar blue assay (Deep Blue Cell Viability^TM^ Kit 424701, Biolegend) was used to analyze cell proliferation at different time points (i.e., day(s) 1, 4, 7 and 14) without disrupting samples, following manufacturer’s instructions. Data were obtained using the Synergy-2 (Biotek) plate reader (Ex 530-570 nm / Em 590-620 nm). All intensity measurements at days 4, 7 and 14 were normalized with their respective reading at day 1. Data were acquired in triplicates on three independent biological experiments.

#### 2.1.3 Biomarkers expression: flow cytometry

Marker expression (i.e., CD44, CD44v6, CD24, E-cadherin, vimentin, ALDH) was analyzed by flow cytometry (BD Fortessa X-20) and following the procedure reported in ^15^. To exclude dead cells from measurements, cells were incubated with 1 μg/mL of 4′,6-diamidino-2-phenylindole (DAPI) solution in 1 × PBS (5 min, RT), then washed with 1 × PBS and re-suspended in 1 × PBS for further measurements. Data were analyzed with FlowJo software (v10.8.0, BD) and for gate single live cells and obtain measurements on median fluorescence intensity (MFI) and number of cells positive for each marker (%). The gates for ALDH+ (%) cells were sorted based on the DEAB negative control of the respective sample. The median intensity of the marker was normalized by its respective isotype control for every sample and then plotted as an average of N=3 independent experiments.

### 2.2 A data-driven approach toward profiling breast cancer phenotypes

The available data were initially grouped based on the TMEs characteristics (**Table 1**) and preliminarily analyzed by computing their descriptive statistics, i.e., the median, the 25^th^ and 75^th^ percentiles of the available records, and the minimum (0^th^ percentile) and maximum (100^th^ percentile) records excluding outliers (**Figure 6B**) for a visual representation of the former statistics through boxplots per biological readouts and TME). Two different data-driven paths are then followed to use the measured markers to characterize inter-phenotype differences and intra-phenotype behaviors, whose main tools and features are described next.

#### 2.2.1 Data processing tools

To unveil the peculiarities of the two considered cell phenotypes with respect to features of the microenvironment, we jointly used a set of off-the-shelf tools from statistics and machine learning, namely the sample Pearson correlation coefficient, k-means ^21^, the silhouette index ^22^ and the mutual information score ^23^. Toward formally introducing these tools, we referred to the *j*-th marker as *x*^*j*^, with *j* ∈ {1, … , *n*} ⊂ *N* being the number of measured markers throughout our experiments, while we indicated the *k*-th feature of the microenvironment as *u*^*k*^, for *k* ∈ {1, … , *m*} ⊂ *N*. Meanwhile, we referred to the *i*-th sample of the *j*-th marker available in our dataset as 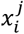, for *i* = 1, … , *N* (with *N* ∈ *N* being the cardinality of the available datasets) and with *j* ∈ {1, … , *n*}. The markers and experimental conditions in each record of our dataset are collected into two vectors, respectively denoted as *x*_*i*_ ∈ *R*^*n*^ (and often referred to as the *feature vector*) and *u*_*i*_ ∈ *R*^*m*^, for *i* = 1, … , *N*.

According to the previous definitions, the Pearson correlation coefficient was used to quantify the strength and direction of the (linear) relations between two markers or a marker and a microenvironment condition based on the available measured markers. Formally, when looking at markers’ correlations, such score can be defined as described in **Equation 1**.

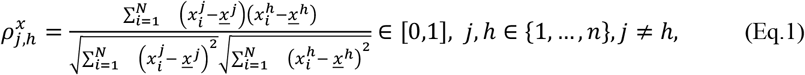

where *<u>x</u>*^*j*^ and *<u>x</u>*^*h*^ are respectively defined as 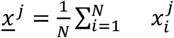 and 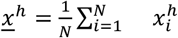, with *j, h* ∈ {1, … , *n*}, *j* ≠ *h*.

The indicator is instead defined in **Equation 2**.

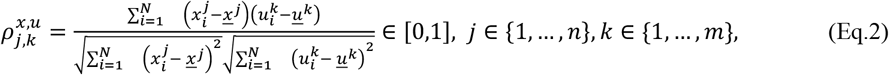

when evaluating the correlation between markers and microenvironment features, with 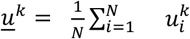 , for *k* ∈ {1, … , *m*}. In this work, these indicators were used to disclose correlations between the measured markers as well as that between them and the microenvironment features when looking at the single cell phenotypes.

Meanwhile, k-means was used to group the available markers into a predefined number of clusters *G* ∈ *N* by looking at central trends in the data, toward discovering the ones that were relevant for distinguishing between phenotypes and the existence of different intra-cluster behaviors. Starting from an initial guess of the groups’ means 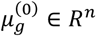 at the 0-th iteration, for *g* = 1, … , *G*, k-mean unfolds by recursively carrying out two steps. At each iteration *q* ≥ 0, with *q* ∈ *N*, each record was first assigned to a cluster based on its Euclidean distance from the current mean, i.e., each group was constructed according to the logic described in **Equation 3**.

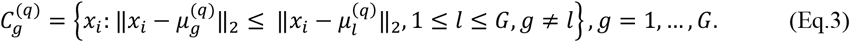

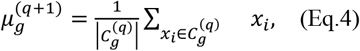

Once the groups were created, the clusters’ means (also called *centroids*) were then updated as described in **Equation 4**, where 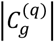 denotes the cardinality of the *g*-th cluster at the *q*-th iteration. This iterative procedure was carried out in an *unsupervised* way (i.e., without requiring the data points to be pre-assigned) until a termination criterion was satisfied, e.g., points’ assignments do not change over two consecutive iterations. Note that, according to the grouping logic in Equation 3, ambiguous points (namely those that can be assigned to multiple clusters) were assigned to one cluster only.

Despite the intuitiveness of k-means, this algorithm rests on the assumption that the number of groups *G* into which the available features have to be partitioned is pre-fixed. While in our setting this is true when distinguishing between cells phenotypes, this was not the case when trying to unveil different behaviors within the same cell line. To determine the number of clusters in this second scenario, we used the silhouette index in **Equation 5**, where *C*_*g*_ is the *g*-th cluster after the termination of k-means and s(xi) is described in **Equation 6**.

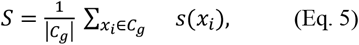

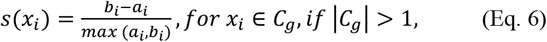

with *a*_*i*_ and *b*_*i*_ respectively used to assess a cluster’s cohesion and its separation from the others, defined as

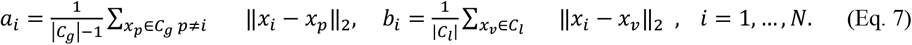

Note that, the silhouette index *S* in Equation 5 ultimately satisfied *S* ∈ [−1,1], with values of *S* close to 1 indicating that samples were coherent and well-separated while *S* close to 0 being negative denoting possible overlaps between clusters or a non-negligible misclassification rate, respectively. Note that, apart from selecting the number of behaviours recognizable from the available markers, in our approach we further used the silhouette index as an indicator of the clusters’ quality, ultimately guiding the choice of the markers that allowed the discrimination between cell phenotypes. Apart from using the silhouette index, the selection of relevant markers for the characterization of cell phenotypes was also performed by using the mutual information score, estimating the mutual information between each marker and the grouping labels available when distinguishing between cell phenotypes and not used by k-means for clustering.

#### 2.2.2 A bird’s eye view on intra-phenotype relationships and inter-phenotype differences

As a preliminary step to understand links between markers, microenvironment conditions and cell phenotypes, our data-driven pipeline consisted of a preliminary step where the Pearson’s correlation coefficients between markers (see Equation 1), as well as that with the different features of the microenvironment, were estimated for each cell line separately. By inspecting them, these coefficients provided initial insights into differences in the relations between markers and microenvironment conditions among different cell lines, while indicating which markers were more likely to react to changes in a specific microenvironment characteristic when considering a single cell line. Nonetheless, looking at Pearson’s correlation was not sufficient to have a quantitative and concrete approach to simplify the experimental scheme and determine the factors that were necessary to differentiate between metastatic and recurrent features of MDA-MB231 compared to the non-recurrent ones of MCF-7.

#### 2.2.3 Are there any discernible intra-phenotype behaviors depending on the microenvironment condition?

Apart from distinguishing among cell phenotypes, further information on intra-phenotype behaviors driven by the different microenvironment conditions can be key to simplifying experiment design (as well as making it more sustainable) while enabling a deeper understanding of the interactions between the microenvironment and the cells. To this end, k-means is exploited to determine whether distinct behaviors could be detected within each cell line depending on microenvironment conditions, using the measured markers as features for clustering purposes. In this case the number of clusters cannot however be selected a priori. Therefore, in detecting behaviors within the same group of cells, we try to separate the markers into an increasing number of clusters up to a maximum of clusters, exploiting the silhouette index (see Equation 5) to evaluate the quality of separation and then decide the amount of different behaviors (if any) detectable within a single cell line. In particular, the number of clusters is ultimately chosen as the one leading to the number of groups that causes the least drop of the silhouette score, picking the minimal number of groups if two tested values for leads comparable silhouette scores. This design choice is made to search for a trade-off between accuracy (i.e., limited drops on the attained silhouette index’s values) and number of identified behaviors. Note that, in this case, no pre-existing labels are available as ground truth to characterize intra-phenotypes behaviors and, thus, feature selection (i.e., unveiling unneeded markers) can only be performed by trial and error, removing one feature at a time and evaluating changes in the resulting silhouette score.

#### 2.2.4 Toward sustainable profiling of the invasive potential of breast cancer cells

As already mentioned previously, the two cell lines considered in this study (MCF-7 and MDA-MB231) are associated with different invasive potentials. Understanding which experimental conditions (see **Table 1**) and measured markers (**Table 2**) allowed one to distinguish between the two cell lines would be key to informing future experiments and guide the analysis of other cell phenotypes. To this end, by using the techniques introduced in Section 2.2.1, we investigated whether all the measured markers (**Table 2**) were all needed to distinguish between the responses of MCF-7 and MDA-MB231, and if there existed also a relationship between the TMEs and the relative importance of the markers in performing such a distinction. Toward grasping such an understanding, we proposed the use of the data-driven pipeline, as schematized in **Figure 4**, and involving three main stages. Initially, k-means (see Section 2.2.1) was used to cluster the available features (i.e., the recorded markers) into two groups by fixing all microenvironment conditions but one. The silhouette index (see **Equation 5**) was then employed to evaluate the separability between the obtained group, hence assessing the quality of the results achieved via the first unsupervised clustering routine. Note that this first phase did not de facto exploit our prior knowledge of the actual cell line (i.e., the *label*) each data record was associated with, in turn allowing us to use all the available (yet scarce from a ML perspective) samples to carry out k-means. Nonetheless, the knowledge of the actual cell line can already be used to check the quality of the grouping one blindly obtains by employing this clustering technique using all the available markers, already giving an insight into whether they were all needed to distinguish between MCF-7 and MDA-MB231. Our prior knowledge of the actual cell line each set of markers is associated with is instead used in the second step of our procedure, i.e., feature selection. To this end, an estimate of the mutual information between each available marker and the actual label (i.e., invasive/non-invasive cell phenotype) was employed to rank the available features, toward unveiling those markers that are likely unnecessary to distinguish between cell lines. This additional step allows for discarding non-informative markers by thresholding the mutual information. The resulting reduced set of features is ultimately clustered again through K-means, and its outcome is validated quantitatively by looking at the silhouette index and comparing the label attributed to the data in an unsupervised fashion with the actual ones. Note that, to grasp the implication of the different microenvironment conditions (i.e., PH, perfusion, matrix stiffness, and composition), this procedure is carried out by fixing one microenvironment condition at a time and not explicitly considering any of the remaining microenvironment characteristics among the features to be clustered via k-means.

## 3. Results and discussion

### 3.1 Intra-phenotype descriptive statistics and Correlation plots

To visualize the correlations among all the parameters of the engineered in-vitro system, we firstly plotted the measured data of cellular phenotypes and the microenvironmental conditions to create two separate Pearson correlation matrices, one for each cell line (**Figure 2**). The complexity of the plot indicates that there are numerous factors to consider when comparing among the two cell lines. Notably, there are significant differences as well as similarities in the correlation patterns among MDA-MB 231 and MCF-7 cells. For instance, in MCF-7 cells, E-cad (both median fluorescence intensity and +ve %) is positively correlated with pH and perfusion, whereas in MDA-MB 231, E-cad is positively correlated with mean hydrogel stiffness. In both the cell lines with respect to B-CSC marker expression, ALDH+ (epithelial type CSCs) is negatively correlated to increase in perfusion and pH, but is positively correlated to stiffness. On the other hand, CD44+/CD24-(mesenchymal type CSCs) is always positively correlated to perfusion. This goes in line with previous in-vivo studies that mention CD44+/CD24-cells are present on the tumor boundary, which is known for high perfusion rates, whereas ALDH+ cells are present in the tumor core known to have a low pH microenvironment ^19,24^. These results reinforce the importance of physico-chemical microenvironment in possible spatial distribution of E-CSCs in the tumor core as compared to M-CSCs at the tumor boundary. Within the same cell line, the varied influence of biophysical parameters is better compared using the correlation plot. For MCF-7, pH and perfusion show a greater correlation with changes (both positive and negative) in cellular phenotypes than hydrogel properties. In MDA-MB 231, most parameters affect cellular phenotypes. However, in both cell lines, hydrogel gelatin content shows little to no correlation with phenotypic changes compared to hydrogel stiffness, implicating that this parameter could be removed in future experimentation.

**Figure 2.**
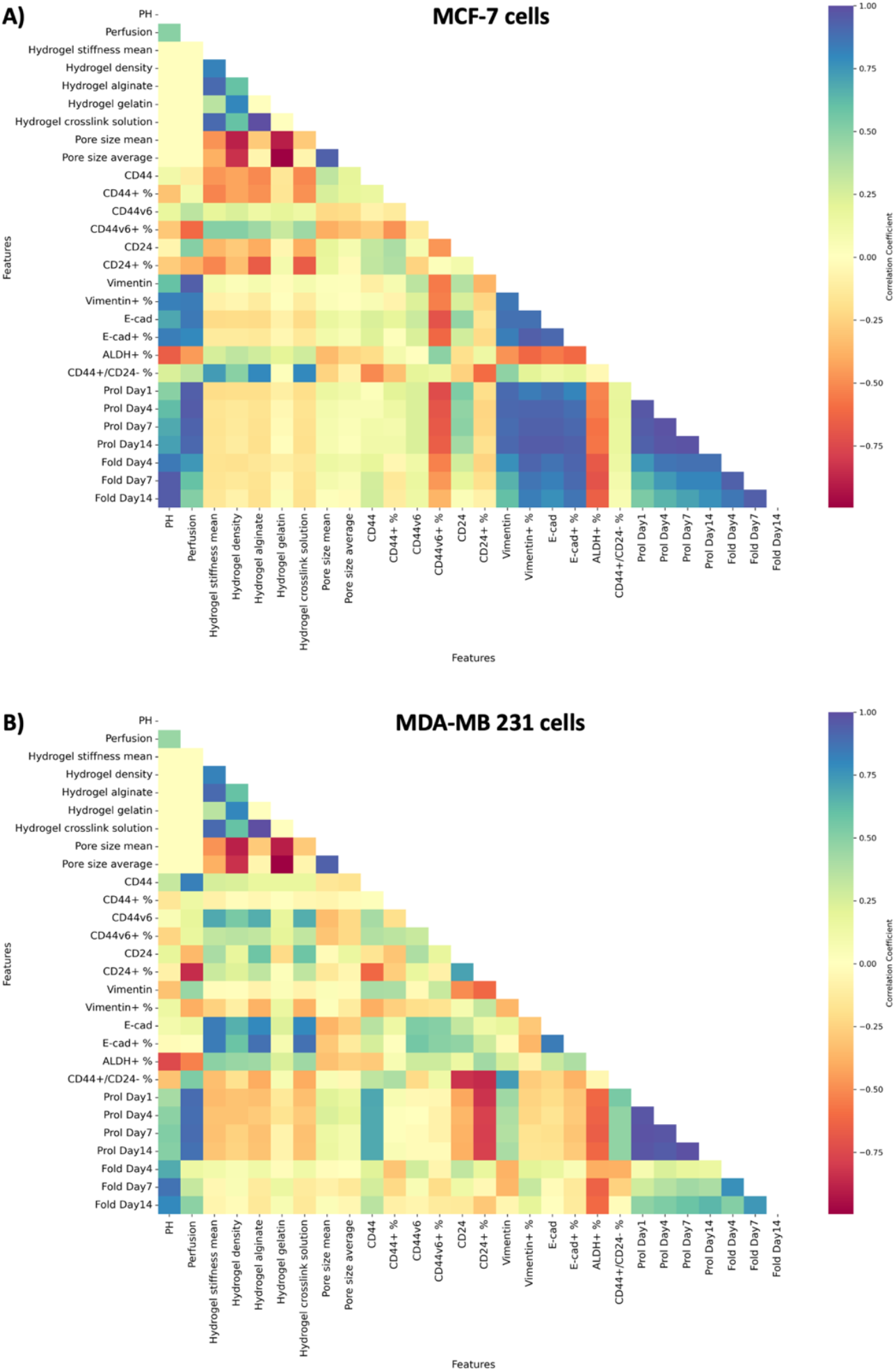
Correlation plot of all features of breast cancer MPS model. Pearson correlation coefficients computed among microenvironmental conditions of the MPS model and measured cellular markers, plotting possible linear correlations of either positive (blue), neutral (yellow) or negative (red) correlation in **A)** non-invasive MCF-7 cells and **B)** invasive MDA-MB231 cells.

In summary, the correlation plot may help in providing a bird’s eye view of the data for visual comparison. It helped in discerning certain patterns with marker expression compared among the two cell lines. In terms of experimental design, the correlation plot aided in visualizing certain biophysical parameters that could be influential in bringing about changes in cellular phenotype. However, it doesn’t provide a robust, quantitative and concrete approach to simplify the experimental scheme and determine which factors would be absolutely necessary to differentiate among the metastatic and recurrent features of MDA-MB231 compared to the non-recurrent ones of MCF-7.

Next, to recognize patterns of cell behavior in differing microenvironments, the cellular outputs or features (proliferation, EMT and B-CSC marker expression) from all microenvironments were clustered for each cell line using k-means. The optimal number of clusters was decided based on silhouette index where the clustering with highest silhouette score was chosen (**Figure SI.1**). As evident for both MDA-MB231 and MCF-7 cell lines the highest silhouette index was when k=2 clusters were formed. To visualize this clustering, among the 16 measured features we firstly plotted three features namely proliferation (day 7), CD44+/CD24- and ALDH+ in different microenvironments where each data point was colored based on their classification in two clusters (as determined previously with all the features) (**Figure 3**).

**Figure 3.**
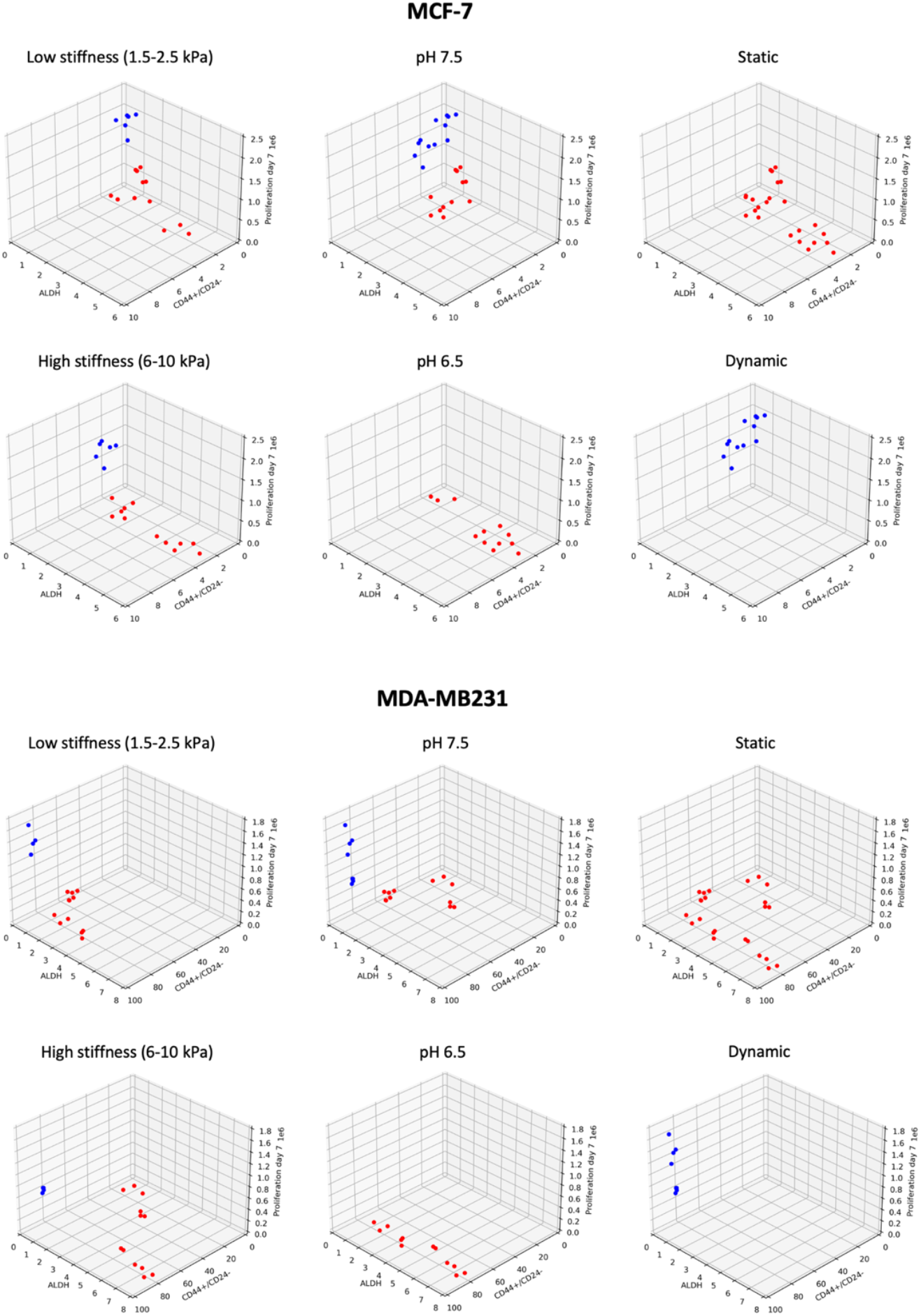
k-means based clustering of cellular phenotypes in different microenvironments. Dot-plots depicting ALDH+ (%), CD44+/CD24-(%) and proliferation (Day 7) in low / high stiffness, pH 6.5 / pH 7.4 and Static / dynamic flow conditions for MCF-7 cells (top) and MDA-MB231 (bottom). Red and blue depict two cluster population based on k-means clustering of all cellular phenotypes in different microenvironments.

**Figure 4.**
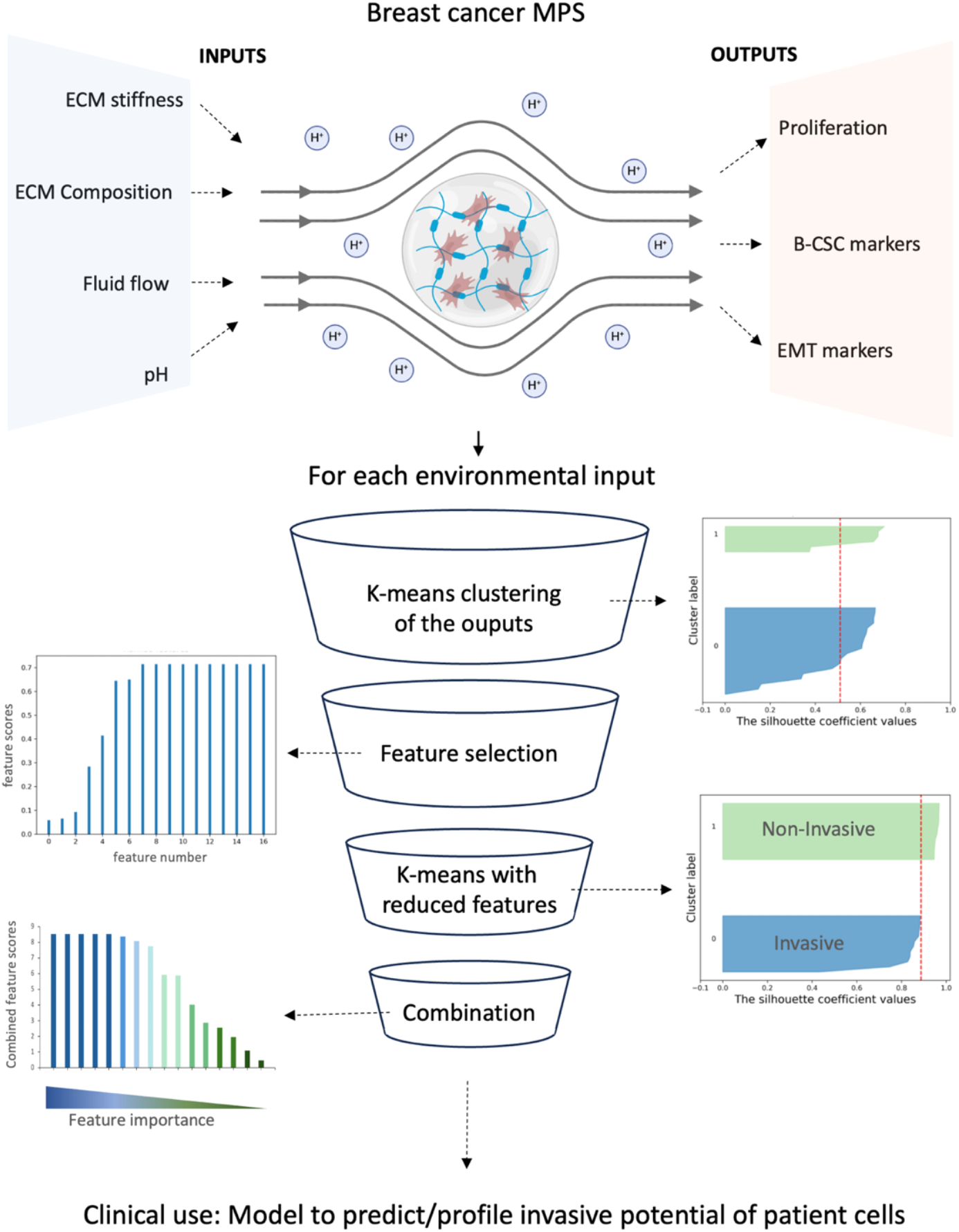
Schematic representation of the digital tools used profiling invasive and recurrent behaviors in breast cancer cells. ML based data pipeline outlining steps to profile invasive MDA-MB231 and non-invasive MCF-7 cells by: 1) Pooling all cellular phenotypes of both cell lines in each microenvironmental condition, 2) Performing unsupervised k-means clustering and keeping the number of cluster constant (K=2), 3) Ranking features/cellular phenotypes based on the alignment of the features with the true label, 4) Performing k-means again with selected and reduced set of features and confirming with silhouette index to plot clusters with true labels, and 5) Combining feature scores from all microenvironments to rank the most important features useful in distinction of invasive phenotypes.

Based on the clusters formed, it is evident that cell lines behave similarly in some microenvironments while different in others. For instance, single clusters are present when cell lines are cultured in pH 6.5, static and dynamic conditions, suggesting these conditions create mostly homogenous population with these cellular phenotypes. For example, dynamic conditions create higher proliferating and low ALDH+ populations whereas pH 6.5 creates low proliferating and high ALDH+ populations. In fact, a tumor core that is characterized by low pH is observed to have presence of high ALDH+ (E-CSCs) populations in vivo ^19^, which is also apparent in our analysis irrespective of the cell line. On the other hand, pH 6.5 and dynamic populations maintain CD44+/CD24-at higher levels in MDA-MB231 and not in MCF-7, creating a distinction in their behavior. This analysis suggests that for streamlining future experiments where we can differentiate between MCF-7 and MDA-MB231, a combination of ‘microenvironment’ and ‘cellular phenotypes’ will have to be considered. However, with these graphs, it is difficult to visualize more than three parameters at a time, which leads to an incomplete analysis. At this stage, dimensionality reduction methods may help in visualization, but will lead to a loss of interpretability, making it challenging to understanding which features differentiate between MDA-MB231 and MCF-7 profiles and defeating the purpose of data-informed experiment streamlining. Overall, this analysis, whilst providing some insights in cell behavior, still fails to provide a robust and scalable method that can be easily adopted for other bioengineering setups or translated to larger clinical datasets.

### 3.2 Data processing pipelines towards profiling the breast cancer invasive potential

To develop a quantitative approach that could inform on features important to differentiate between MDA-MB 231 and MCF-7 profiles, we utilized a data processing pipeline based on unsupervised k-means clustering, feature extraction and feature selection was applied with all the parameters of the MPS model (**Figure 4**). However, for this method, cellular data from both the cell lines was used together for clustering. Briefly, output parameters, representing distinct cellular phenotypes, were clustered for each microenvironmental input parameters, with silhouette indexes used to assess the initial unsupervised cluster separation. By comparing to actual labels, cellular phenotypes were ranked based on their contribution in distinguishing MDA-MB 231 and MCF-7 profiles. Eventually low-ranked features were removed, and clustering was re-evaluated to validate differentiation accuracy (**Figure 4, Figure 5**).

**Figure 5.**
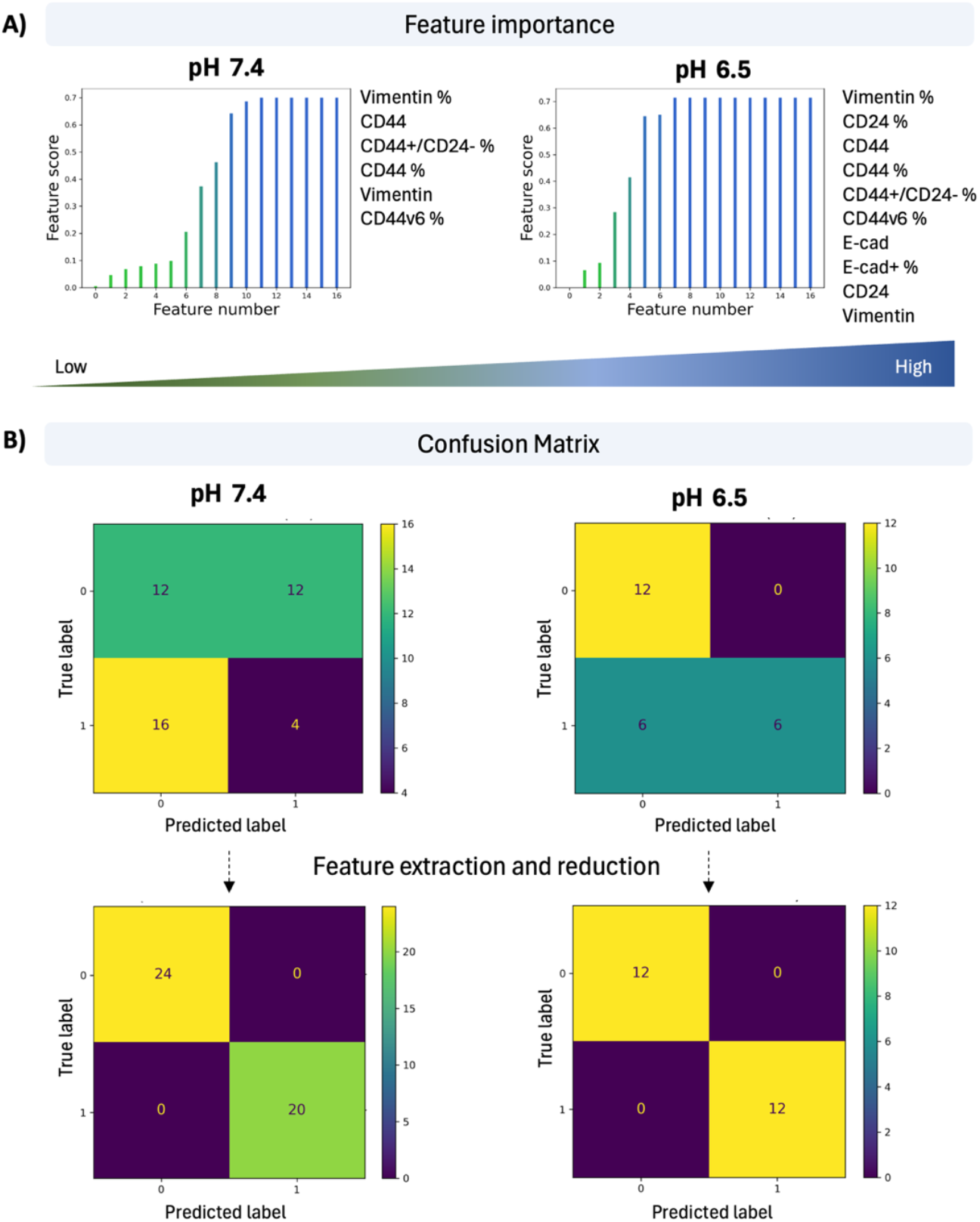
Ranked feature scores of cellular phenotypes in a microenvironmental input. **A)**Example of feature scores plot where features/cellular phenotypes are ranked from low to high (left) and the list of most important features (right) when BC cells are cultured in pH 7.4 and pH 6.5. **B)** Confusion matrix showcasing true positive (upper left), false negative (lower left), false positive (upper right) and true negative labels (lower right) before and after feature extraction in pH 7.4 and pH 6.5 conditions.

**Figure 6.**
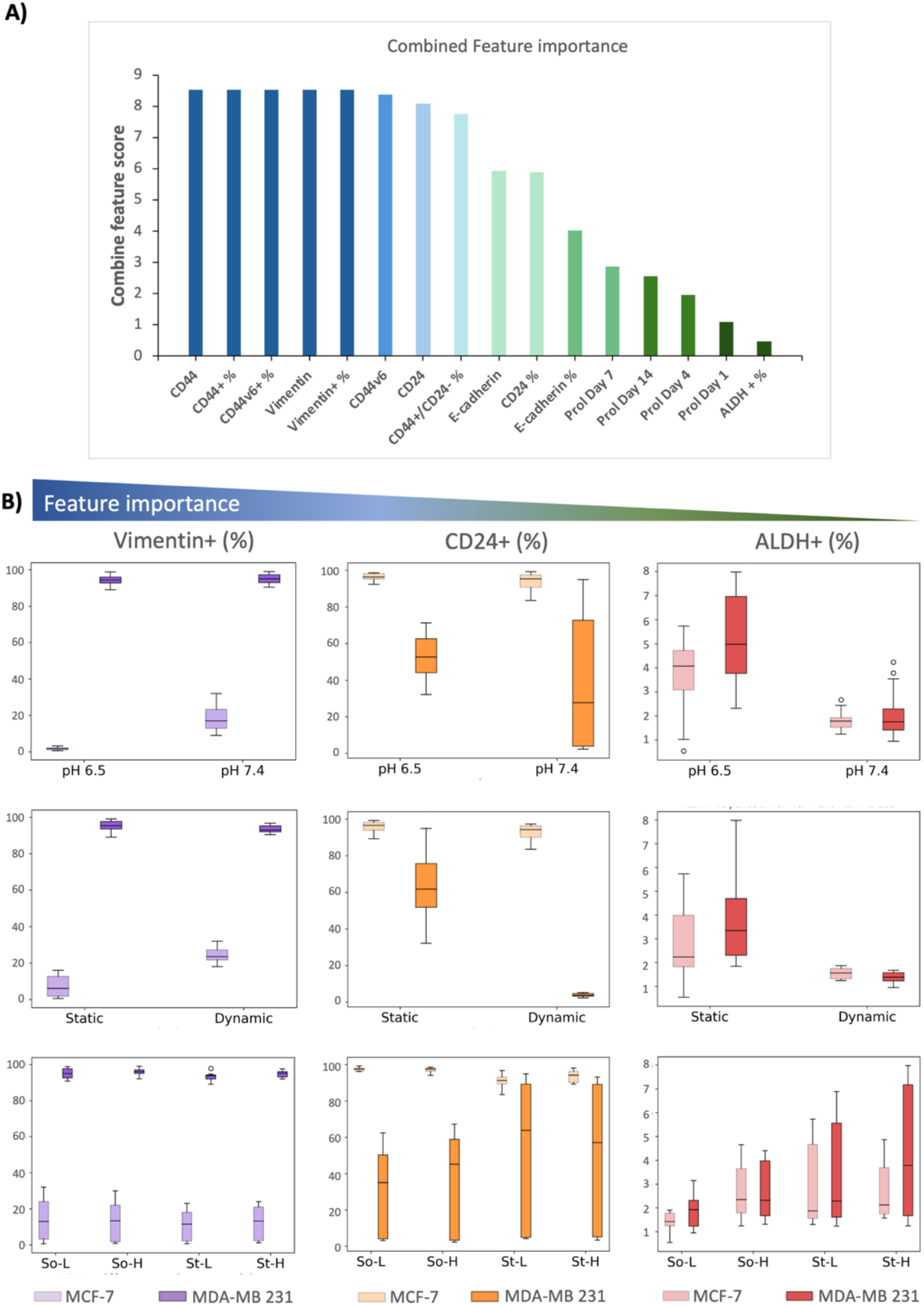
Combined feature importance of cellular phenotypes. **A)** Combined feature score of the cellular phenotypes in all the microenvironments, arranged from the highest to the lowest. **B)** Bar plot representation of Vimentin (%) - high importance, CD24 (%) - intermediate importance and ALDH (%) - low importance in different microenvironments for non-invasive MCF-7 (light shade) and invasive MDA-MB 231 (darker shade).

By using this method, we were able to extract features that were absolutely necessary to distinguish MDA-MB231 vs MCF-7 in each microenvironmental condition. The output/features from each microenvironmental condition namely the pH (pH 7.4 or pH 6.4) (**Figure 5**), perfusion status (static or dynamic), matrix stiffness (hydrogels So-L, So-H, St-L, St-H) or matrix composition (Alginate or gelatin content) were processed with the pipeline and the features were ranked (**Supplementary Information, Figure SI.2, Table SI.1**). Interestingly, each microenvironment had a different set of features that were important for distinction between the two cell lines (**Figure SI.2**). For example, for pH 6.5 dynamic perfusion and soft matrixes resulted in the highest number of important features necessary for this distinction, suggesting that these microenvironments may help in better isolating the invasive phenotypes from non-invasive ones. A confusion matrix table that showcases how well a machine learning model performs by comparing predicted values to actual values in a dataset, was used to assess the accuracy of the ML classification model after feature reduction for each condition (**Figure 5**). As confirmed by this matrix, both false positives and false negatives become zero after feature extraction and reduction. Overall, this data pipeline streamlines the experimental process and aids in identifying which features distinguish between invasive and non-invasive profiles if a specific microenvironmental condition is selected.

### 3.3 Unveiling the markers needed to detect invasive cell phenotypes

To determine which features were consistently ranked as important across all microenvironmental conditions, we aggregated the feature scores of all the outputs represented in **Supplementary table SI.1**. The features that frequently scored highest were CD44, CD44+(%), CD44v6, CD44v6(%), Vimentin, and Vimentin+(%), while the lowest-scoring features were cellular proliferation and ALDH+(%) (**Figure 6**). Histograms revealed that the expression of high-importance features like Vimentin was consistently lower in MCF-7 or non-invasive profiles, even under tumorigenic conditions such as low pH, high stiffness, and dynamic environments, whereas in MDA-MB231, Vimentin+ population was always high. For features of moderate importance, such as CD24+, MDA-MB231 displayed variable levels depending on the microenvironment, sometimes indistinguishable from MCF-7. Low-importance features like ALDH+ showed lower levels in both cell lines, with no significant differences among them across all microenvironments. The histogram/box-plot analysis supports the conclusions from our data pipeline, which suggests that this approach can provide a robust quantitative methodology for feature selection and develop a data-driven sustainable approach to narrow down experimental parameters. It was observed that mesenchymal markers and the CD44+/CD24-status were more important than the epithelial marker (E-cadherin) and ALDH+ status. This indicates that within the CSCs population, mesenchymal CSCs (M-CSCs) are better predictors of a recurrent phenotype than epithelial CSCs (E-CSCs).

### 3.4 Combination of breast-specific MPS and ML models as new predictive tool

MPS models integrate human cells with engineered microenvironments to mimic key physiological responses, providing a more dynamic and realistic setting than conventional cell culture systems. MPS aims to simulate the mechanical, chemical, and biological stimuli that tissues experience *in vivo*, making them powerful tools for drug testing, disease modeling, and personalized medicine ^25–27^. However, the complexity of MPS models lies in their intricate design and functionality. Recreating the microenvironment, ensuring proper cell-cell and cell-matrix interactions, and maintaining fluid flow and other mechanical cues is technically demanding for high throughput experiments. Integrating multiple variables for studying systemic responses can prove a challenge in standardization, scalability, and reproducibility as it leads to complex experimental settings and loss of quick data interpretation. There is no fixed definition for the optimal balance between model complexity and its accuracy or efficacy in delivering relevant in vivo physiology insights for a specific application. This holds true for most in-vitro campaigns where preliminary experiments often include or exclude specific variables arbitrarily, without a rigorous rationale. A more systematic approach to variable selection would enhance the quality and impact of *in vitro* research, fostering more dependable data that can be built upon confidently in other studies.

In this work, a previously used breast cancer MPS model that mimics various physico-chemical aspects of the primary tumor microenvironment was used together with the high dimensional experimental data from various biological, mechanical, and chemical interactions in two cell lines. We selected twelve different microenvironments that mimicked combinations of normal breast tissue (e.g., soft hydrogel, pH 7.4, and static) and tumor breast tissue microenvironments (e.g., stiff hydrogel, pH 6.5 and dynamic). This ensured that we recorded potential cellular phenotypes in changing microenvironments and not just snapshots that are usually measured as markers for disease progression. We measured proliferation, EMT and B-CSC markers as these are all knowingly playing a role in breast cancer progression, especially in metastatic invasion which is the cause of most cancer-related deaths ^28,29^. However, with 12 different microenvironments and 16 phenotypic/marker readouts, it was difficult to compare the consequent multidimensional data and to draw conclusions on which experimental settings brought out distinct behavioral patterns of these cell lines.

ML algorithms can help to cope with high-dimensional datasets efficiently, identifying patterns that may help in optimizing experimental conditions and, thus, tests which are otherwise time-consuming. In most preliminary bioengineering setups, experiments are performed with trial and error, wherein variations are introduced in the *in vitro* model and preliminary read-outs are manually measured. Although the data is not extensive, it still has high dimensions, thus making it difficult to manually analyze them while having a comprehensive understanding of the information embedded in the data. In turn, this makes it harder to use these insights to inform future directions with experimental schemes. In this study, we have explored machine learning pipelines based on an unsupervised approach, k-means, along with feature extraction to inform on the most important features of our breast cancer MPS model that differentiate invasive MDA-MB-231 phenotypes from non-invasive MCF-7. Unlike extensive multi-omics data or patient-centric medical/clinical data ^30^, preliminary bioengineering experiments typically involve less extensive data, preventing the use of more complex techniques due to the risk of overfitting. The proposed procedure allows for evaluating whether different cell phenotypes can be detected by leveraging all the available information, without using dimensionality reduction techniques like Principal Component Analysis (PCA) and Uniform Manifold Approximation and Projection (UMAP), which provide a systematic way to visualize data in a lower dimension space but might hamper the interpretability of the data analysis’ outcomes.

With unsupervised k-means clustering and feature extraction we firstly observed that each microenvironment had a different set of important features that distinguished between the two cell lines. Some microenvironments like acidic pH, dynamic environment and soft matrix created more differentiating features between invasive and non-invasive phenotypes suggesting that these micro environmental conditions can be further used to validate data with other cell lines and patient samples (**Figure 4, Figure 5**). Overall, through this data driven approach, mesenchymal markers such as Vimentin, CD44 and CD44v6 were quantified to be the most important features for the distinguishing cell phenotypes (**Figure 6**). In our experimental setting, this emphasizes that even in differing microenvironments of the primary tumor, M-CSCs could be a better discerning factor for invasion and recurrence than E-CSCs and proliferation. In other words, tumor cells that have the potential to express mesenchymal stem cell markers in various microenvironments could hold the potential to metastasize as well. For this application, the ML model could further be consolidated by training on extensive data from patient cells of known recurrence status in breast MPS models. Compared to cell lines, primary cells can capture patient and sub-type specific variations which may increase the resolution of the prediction made by this model, especially for clinically relevant prognosis such as ‘time to recurrence’.

Thanks to the relative simplicity of the employed tools, the same ‘blue-print’ of data pipeline can be used to investigate and integrate parameters from future experiments as well as other preliminary biology experiments. These can include additional cellular phenotypes as well as patients’ biopsies (either ECM or cells, or both) to translate findings quickly to the clinical setting. With this pipeline we provide a new data-driven approach to look at the data generated by MPS models and a better way to dissect cell-material interaction. This unique combination of bioengineering, tumor biology and ML could further help in creating sustainable experimental schemes for developing prognostic tools and personalized therapeutics.

## 4. Conclusions

Identifying the relationships between biological readouts and the different TMEs, as well as their likelihood to impact cell phenotype, is crucial when using patient-derived cells to predict therapeutical outcomes through *in vitro* experiments. This information is key for the design of specific MPSs and sustainable experimental campaigns focusing on the identification of relevant experimental conditions that not only result in time and resource savings but mainly increase the impact of the research.

The novel combination of engineered breast-specific MPSs and ML models proposed in this work represents a first step toward a data-guided outlook on cell/material interactions, which may help in understanding the interplay between TME features and cancer-related biological onsets. The quantitative indications on the TMEs impact on cancer cells behavior provided by our approach can ultimately steer future experiments, making *in vitro* approaches more sustainable. At the same time, such an understanding may be helpful at a clinical level, guiding the drug design of cancer therapies. Due to the straightforward nature of the tools used, the same data pipeline framework can be applied to analyze and incorporate parameters from future clinical experiments and/or early-stage biological studies to facilitate the rapid translation of findings into clinical applications.

## Notes

### Competing Interest Statement

The authors have declared no competing interest.

