## Supplemental figures and tables for "A data-informed approach for engineering in-vitro experiment design to decipher key features of invasive breast cancer cell phenotypes"

[1] Biotech/DII-UNITN

[2] TU/e

### SUPPLEMENTARY INFORMATION

### SI.1 Silhouette plots

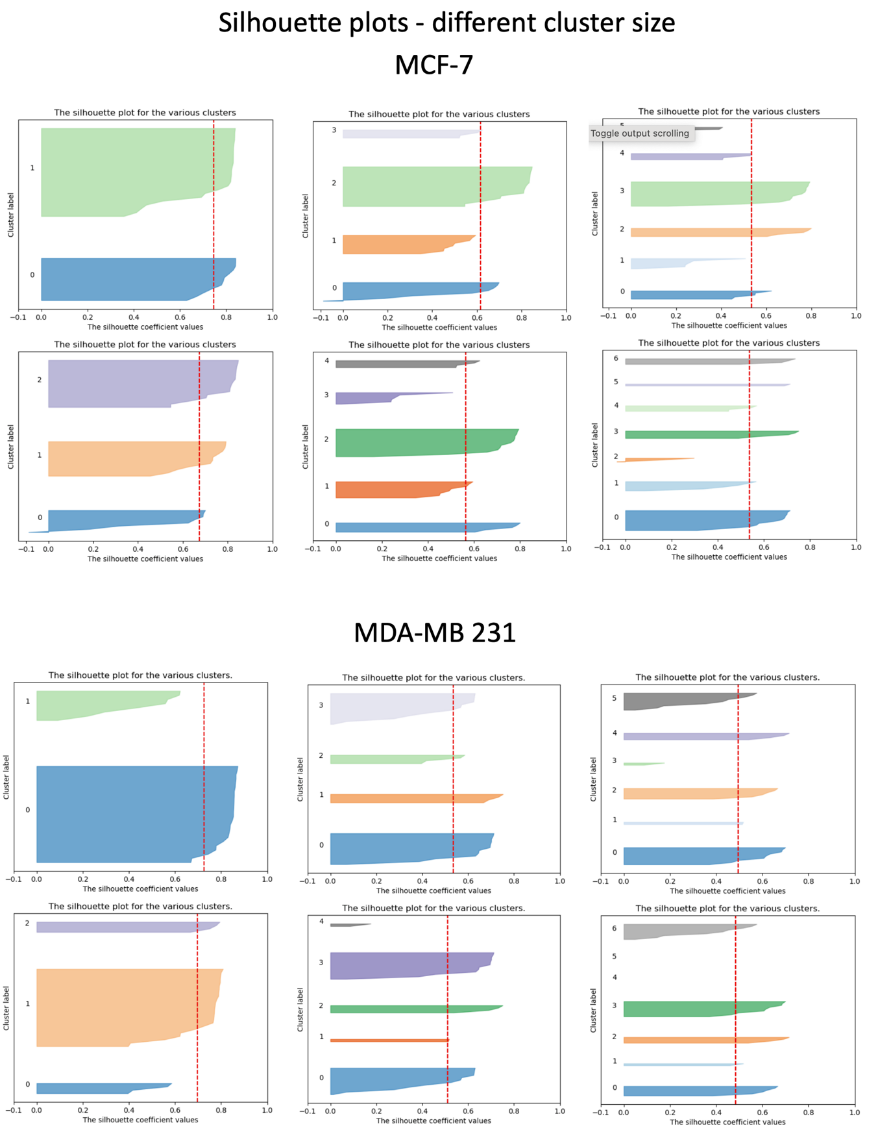

**Figure SI.1. Silhouette plots:** Silhouette plots of retrieved cellular phenotypes (from all microenvironment) when different cluster sizes (k=2 to k=7) are chosen for MCF-7 cells (top) and MDA-MB231 cells (bottom)

**Commented [AT1]:** Missing figure caption

### SI.2 Feature importance of cell phenotypes in all microenvironments

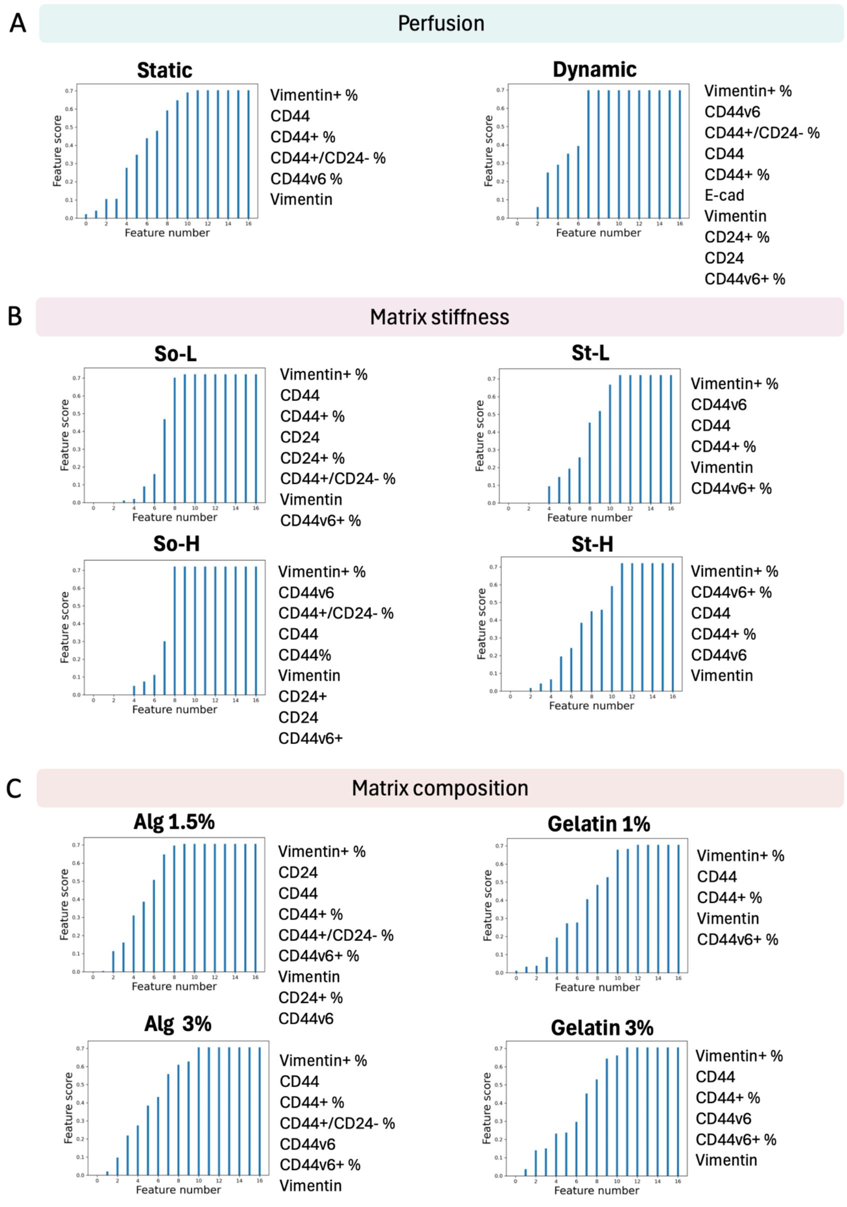

**Figure SI.2. Feature importance:** Cellular phenotypes/ features ranked from lowest to highest based on their importance in distinguishing MCF-7 vs MDA-MB231 in varying **A)** Perfusion status, **B)** Matrix stiffness and **C)** Matrix composition. The highest ranking features with equal importance score are listed on the right of each graph.

**SI.3 Ranked features**

**Table SI.1.** List of ranked features/markers in each microenvironment, ranked lowest to highest based on their importance in distinguishing invasive features

| **pH 6.5** | **pH 7.4** | **Static** | **Dynamic** | **So-L 1.8** | **So-H 2.4** | **St-L 6.1** | **St-H 10.1** | **Alg 1.5%** | **Alg 3%** | **Gel 1%** | **Gel 3%** |
|---|---|---|---|---|---|---|---|---|---|---|---|
| ALDH | ALDH | Prol Day1 | E-cad+ (%) | Prol Day1 | ALDH | Prol Day1 | Prol Day4 | ALDH | ALDH | ALDH | ALDH |
| Prol Day1 | Prol Day4 | ALDH | ALDH | Prol Day7 | Prol Day7 | ALDH | ALDH | Prol Day1 | alginate | Prol Day1 | Prol Day4 |
| Prol Day4 | Prol Day14 | Prol Day4 | Prol Day14 | Prol Day4 | Prol Day14 | Prol Day14 | Prol Day7 | E-cad+ (%) | Prol Day4 | Prol Day14 | Prol Day1 |
| Prol Day7 | E-cad+ (%) | Prol Day7 | Prol Day1 | Prol Day14 | Prol Day4 | CD24+ (%) | Prol Day14 | Prol Day4 | CD24+ (%) | Prol Day4 | Prol Day7 |
| CD44v6 | Prol Day1 | Prol Day14 | Prol Day4 | ALDH | Prol Day1 | Prol Day7 | Prol Day1 | Prol Day14 | Prol Day7 | Prol Day7 | Prol Day14 |
| Prol Day14 | Prol Day7 | CD24+ (%) | Prol Day7 | E-cad+ (%) | E-cad+ (%) | E-cad | CD24+ (%) | Prol Day7 | Prol Day14 | E-cad+ (%) | E-cad+ (%) |
| Vim | CD24+ (%) | E-cad | CD44v6+ (%) | E-cad | E-cad | Prol Day4 | E-cad | E-cad | E-cad | CD24+ (%) | CD24+ (%) |
| CD24 | E-cad | E-cad+ (%) | CD24 | CD44v6 | CD44v6+ (%) | CD44+ / CD24- (%) | E-cad+ (%) | CD44v6 | E-cad+ (%) | E-cad | E-cad |
| E-cad | CD24 | CD24 | CD24+ (%) | CD24 | CD24 | E-cad+ (%) | CD44+ / CD24- (%) | CD24+ (%) | CD24 | CD44+ / CD24- (%) | CD44+ / CD24- (%) |
| E-cad+ (%) | CD44v6 | CD44v6 | Vim | CD24+ (%) | CD24+ (%) | CD24 | CD24 | Vim | Vim | CD24 | CD24 |
| CD44v6+ (%) | CD44v6+ (%) | Vim | E-cad | Vim | Vim | CD44v6+ (%) | Vim | CD44v6+ (%) | CD44v6+ (%) | CD44v6 | Vim |
| CD44+ / CD24- (%) | Vim | CD44v6+ (%) | CD44+ (%) | CD44+ / CD24- (%) | CD44+ (%) | Vim | CD44v6 | CD44+ / CD24- (%) | CD44v6 | CD44v6+ (%) | CD44v6+ (%) |

| | | | | | | | | | | | |
|---|---|---|---|---|---|---|---|---|---|---|---|
| CD44 + (%) | CD44 + (%) | CD44 + / CD24- (%) | CD44 | CD44 + (%) | CD4 4 | CD44 + (%) | CD44 + (%) | CD44 + (%) | CD44 + / CD24- (%) | Vim | CD44 v6 |
| CD44 | CD44 + / CD24- (%) | CD44 + (%) | CD44 + / CD24- (%) | CD44 | CD4 4+/C D24- (%) | CD44 | CD44 | CD44 | CD44 + (%) | CD44 + (%) | CD44 + (%) |
| CD24 + (%) | CD44 | CD44 | CD44 v6 | CD44 v6 + (%) | CD4 4v6 | CD44 v6 | CD44 v6 | CD24 | CD44 | CD44 | CD44 |
| Vim + (%) | Vim + (%) | Vim + (%) | Vim + (%) | Vim + (%) | Vim + (%) | Vim + (%) | Vim + (%) | Vim + (%) | Vim + (%) | Vim + (%) | Vim + (%) |
